# Genomic analysis of *Coccomyxa viridis*, a common low-abundance alga associated with lichen symbioses

**DOI:** 10.1101/2023.09.13.557537

**Authors:** Gulnara Tagirdzhanova, Klara Scharnagl, Xia Yan, Nicholas J. Talbot

## Abstract

Lichen symbiosis is centered around a relationship between a fungus and a photosynthetic microbe, usually a green alga. In addition to their main photosynthetic partner (the photobiont), lichen symbioses can contain additional algae present in low abundance. The biology of these algae and the way they interact with the rest of lichen symbionts remains largely unknown. Here we present the first genome sequence of a non-photobiont lichen-associated alga. *Coccomyxa viridis* was unexpectedly found in 12% of publicly available lichen metagenomes. With few exceptions, members of the *Coccomyxa viridis* clade occur in lichens as non-photobionts, potentially growing in thalli endophytically. The 45.7 Mbp genome of *C. viridis* was assembled into 18 near chromosome-level contigs, making it one of the most contiguous genomic assemblies for any lichen-associated algae. Comparing the *C. viridis* genome to its close relatives revealed the presence of traits associated with the lichen lifestyle. The genome of *C. viridis* provides a new resource for exploring the evolution of the lichen symbiosis, and how symbiotic lifestyles shaped evolution in green algae.

## Introduction

The discovery of symbiosis started with lichens – complex symbiotic assemblages, in which symbiotic partners are tightly integrated into a single body (the thallus), which is often three dimensional and separated into tissue-like layers [1]. The classic definition of a lichen involved two, rarely three, partners: one fungus (the mycobiont), plus one microscopic green alga (the photobiont) and/or one cyanobacterium, which collectively constitute a lichen. This definition, however, has proven too simplistic, since many lichens contain microbial organisms in addition to the main partners, typically bacteria and yeasts [2–5].

Studies on algae in lichens have also shown surprising diversity. Instead of one algal strain, as had been assumed before, some lichens contain two or more coexisting within the same thallus (reviewed by Muggia et al. [6]2018). Often, such co-occurring algae are closely related species of *Trebouxia* or another typical photobiont genus (for examples see [7–11]). Although direct evidence is still lacking, these algae are believed to reside within the algal layer together and occupy more or less the same niche in the symbiotic relationship. In other words, algal cells that were previously assumed to belong to one uniform photobiont, turned out to be two different, albeit closely related photobionts. However, this is not the only instance of algal diversity within lichens.

Compared to lichen photobionts, much less is known about other algae present in lichens. Coming from diverse groups and present in miniscule amounts, these ‘additional’ green algae have recently been termed the phycobiome [12] They are mostly detected in two ways. First, by culturing them from a lichen thallus (e.g., [13–16]). Sometimes this happens by accident, when researchers attempt to culture lichen photobionts [17]. Second, during metabarcoding surveys, in which, when reported, non-photobiont algae represent a small fraction of produced data (e.g., [18–21]). These algae are sometimes assumed to grow epiphytically on lichens (e.g., [13,14]), which is supported by studies comparing algal sequences from washed and unwashed lichen thalli [18,22]. However, much remains unknown about the non-photobiont lichen-associated algae, and the hypothesis that they partake in lichen symbioses cannot be ruled out.

Here, we examine one such alga, *Coccomyxa viridis*, and its distribution in lichen symbioses. We report the first sequenced genome of a non-photobiont lichen-associated alga, and compare it to genomes of closely related algae. We identify characteristics of its genome that are consistent with its lichen-associated lifestyle and discuss its potential role within the symbiosis.

## Results

### Non-photobiont *Coccomyxa* cultured from a *Xanthoria* lichen

A strain of green alga (Fig. 1A) was cultured from a thallus of the *Xanthoria parietina* lichen (Fig 1B). While we originally expected to isolate *Trebouxia*, the main photobiont of *Xanthoria* [23], it appeared to be overgrown by a different alga. Instead of globular cells with a large star-shaped and centrally-located chloroplast, as is the case for *Trebouxia* (Fig. 1A,C, [24]), the algal cells in our cultures were ellipsoidal with chloroplasts in the cell periphery (Fig. 1A).

**Fig. 1.**
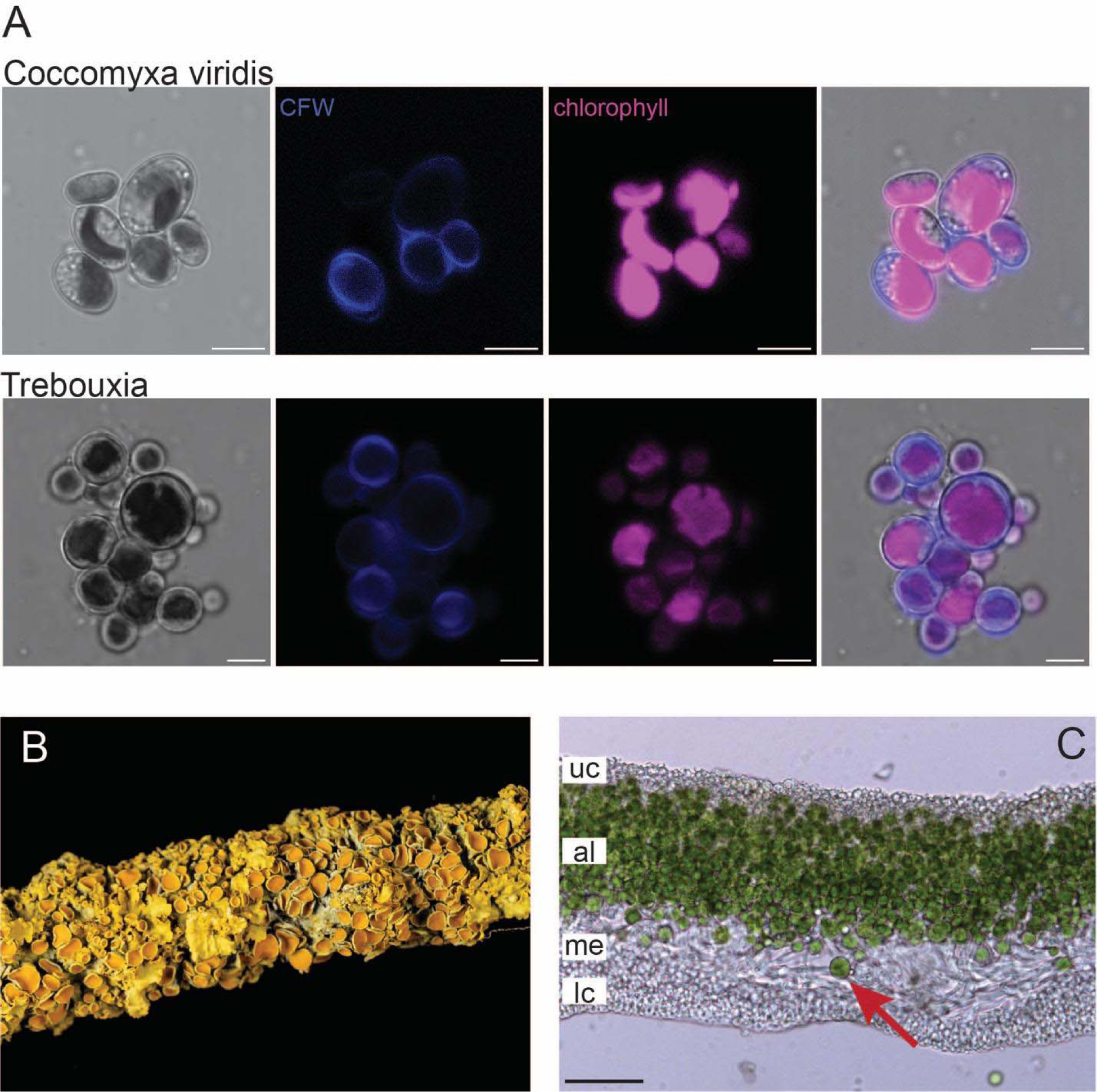
Algae in *Xanthoria parietina.* A. Micrographs of *Coccomyxa viridis* cultured from a *X. parietina* thallus. *C. viridis* cells are ellipsoid and have one or several chloroplasts located near cell exterior. For comparison, the lower track shows the main photobiont of *X. parietina*, *Trebouxia. Trebouxia* cells are globular and contain one centrally located chloroplast, often star-shaped or lobed. In both cultures, cell walls were stained with the Calcofluor White (CFW) stain. Scale bar = 5 μm. B. Thallus of *X. parietina* growing on a tree branch; photo courtesy of Phil Robinson. C. Cross-section through a *X. parietina* thallus, showing internal structure with four layers: uc = upper cortex (formed primarily by mycobiont hyphae embedded in an extracellular matrix), al = algal layer (mycobiont hyphae and photobiont cells), me = medulla (loosely arranged mycobiont hyphae), lc = lower cortex (mycobiont hyphae embedded in an extracellular matrix). The arrow points to a cell of *Trebouxia* residing in the algal layer. Scale bar = 50 μm.

By constructing a phylogenomic tree with publicly available Trebouxiophyceae genomes, we identified our alga as a part of the larger *Elliptochloris* clade (Fig. 2A). An ITS-based phylogeny placed our strain in the *Coccomyxa viridis* clade (Fig. 2B).

**Fig. 2.**
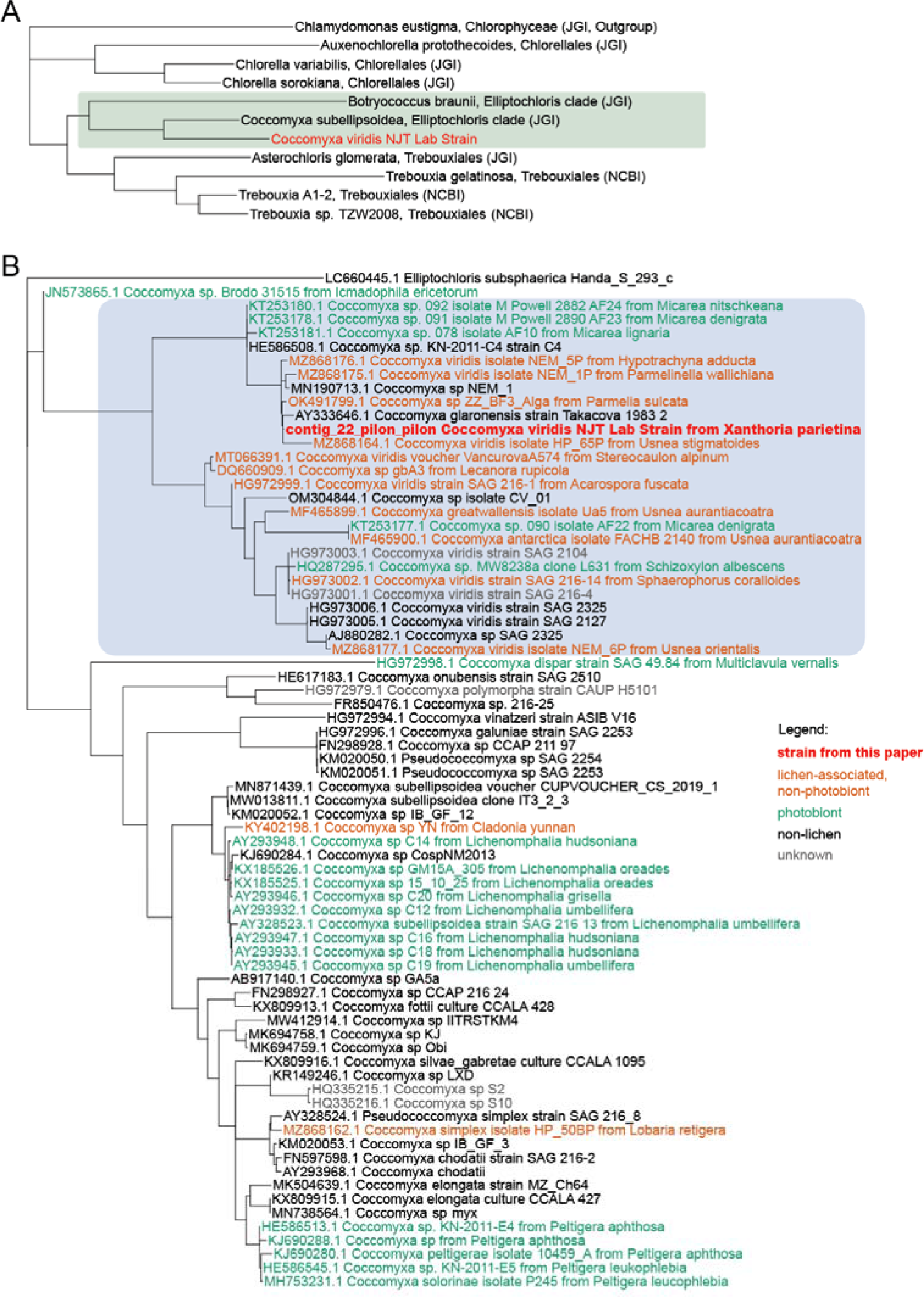
Maximum-likelihood phylogenetic trees showing the taxonomic placement of the studied strain (highlighted in red). A. Phylogenomic tree based on 196 single-copy orthologs. The green rectangle highlights the *Elliptochloris*-clade. B. Phylogeny based on ITS sequences. The blue rectangle highlights the *C. viridis* clade.

### *Coccomyxa viridis* detected in 12% of lichen metagenomes

Apart from identifying the target strain, our phylogenetic analysis showed that the majority of its known close relatives are also lichen-associated. The *C. viridis* clade contained numerous lichen-associated strains, which can be broken into two categories (Fig. 2B). Firstly, five *Coccomyxa* strains that are main photobionts of their lichens: three different *Micarea* lichens and *Schizoxylon albescens*. Of the lichen photobionts included in this analysis, *C. viridis* represented only a minority, as most of them were recovered in a different part of the tree, in the *C. subellipsoidea* and *C.simplex/C. solorinae* clades. Secondly and more notably, 11 non-photobiont alga – i.e. algae cultured from lichens that do not have *Coccomyxa* as their main photobiont. This begged the question of how widespread *C. viridis* is in lichens.

We screened 438 publicly available lichen metagenomes for the presence of *Coccomyxa* ITS. In total, we found 84 *Coccomyxa* sequences. While not all of them came from the *C. viridis* clade, the majority (82%) were identified as *C. viridis* (Fig. 3A). *C. viridis* was present in 53 lichen metagenomes (12% of the screened metagenomes), coming from different lichen groups, collectors, and geographic locations (Fig. 3B-G, Table S1). Nearly all of these algae (88%) are likely not the main photobionts of their respective lichens, as they were detected in lichen symbioses known to have non-*Coccomyxa* photobionts (Table S1).

**Fig. 3.**
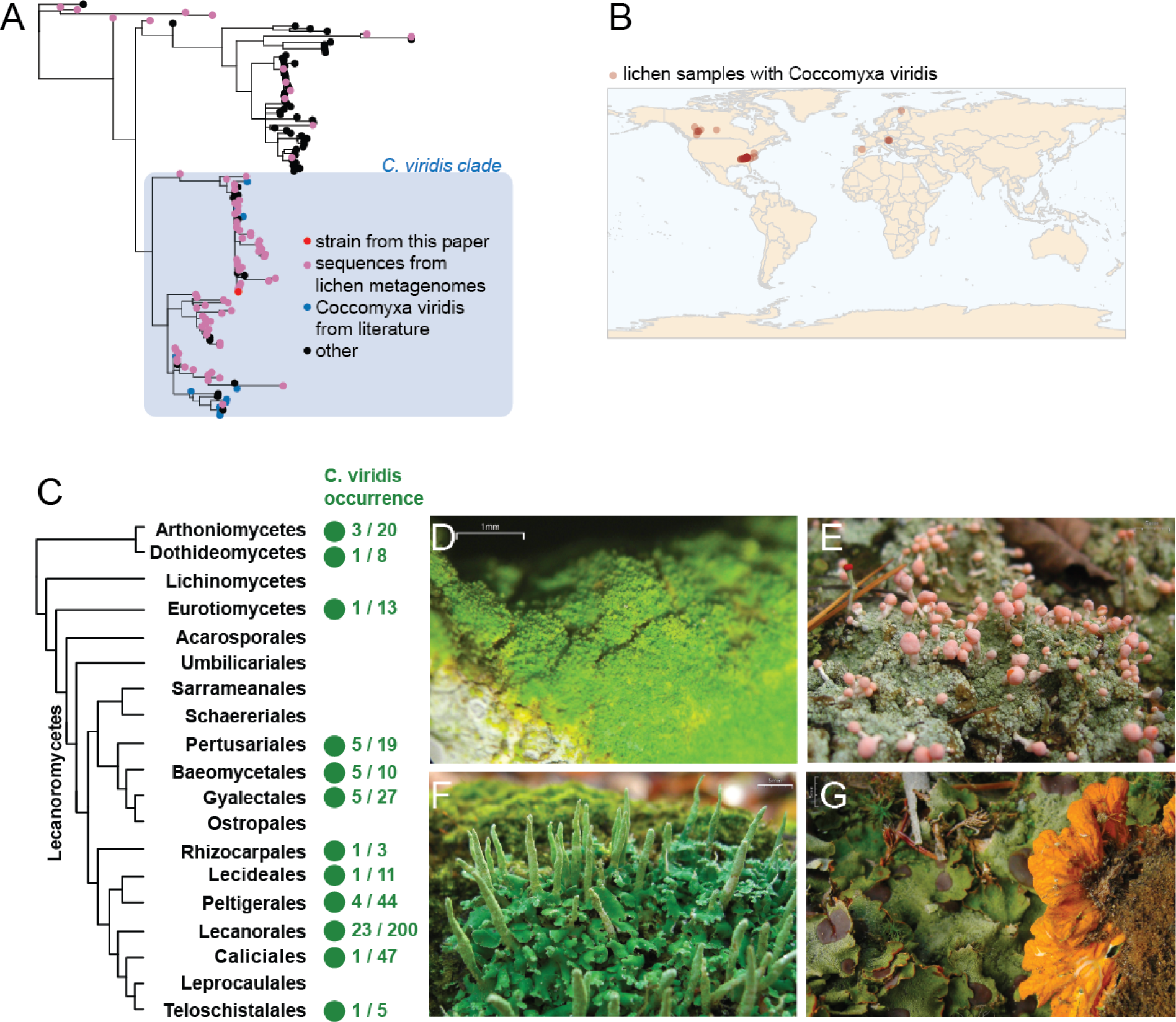
*C. viridis* presence in publicly available lichen metagenomic data. A. Phylogenetic tree of *Coccomyxa* ITS sequences. Pink dots represent sequences pulled from lichen metagenomic data. Blue represents *C. viridis* clade and *C. viridis* sequences from the literature. B. Map showing geographic locations of each lichen sample, in which we detected *C. viridis* by screening metagenomic data produced from this sample. All these samples were collected in North America and Europe, however the real distribution of *C. viridis* could be broader, given that existing metagenomic data on lichens is geographically biased towards these two continents. C. Presence of *C. viridis* across lichen taxonomic groups. The tree represents the phylogeny of lichen mycobionts modified from Tagirdzhanova et al. [67] and Díaz-Escandón et al. [71]; only taxa included in the metagenomic screening are shown. Green dots show taxonomic groups for which *C. viridis* was detected, with the prevalence ratios shown to the right. D-G. Examples of lichen symbioses containing *C. viridis*; photos courtesy of Jason Hollinger. D. *Chrysothrix xanthina* (Arthoniales, Arthoniomycetes). E. *Dibaeis baeomyces* (Pertusariales, Lecanoromycetes). F. *Cladonia ochrochlora* (Lecanorales, Lecanoromycetes). G. *Solorina crocea* (Peltigerales, Lecanoromycetes).

Is *C. viridis* external or internal to the symbiosis?

We washed eight samples of the *X. parietina* lichen and screened the resulting samples via PCR. After both gentle and aggressive washing, three thalli out of four still contained detectable *C. viridis* DNA (Fig. 4, Fig. S1). In contrast, only one wash water sample had traces of *C. viridis*. Generic algal primers yielded *Trebouxia* sequences for all samples, confirming that *Trebouxia* and not *C. viridis* is the main photobiont of these lichens. (Table S2).

**Fig. 4.**
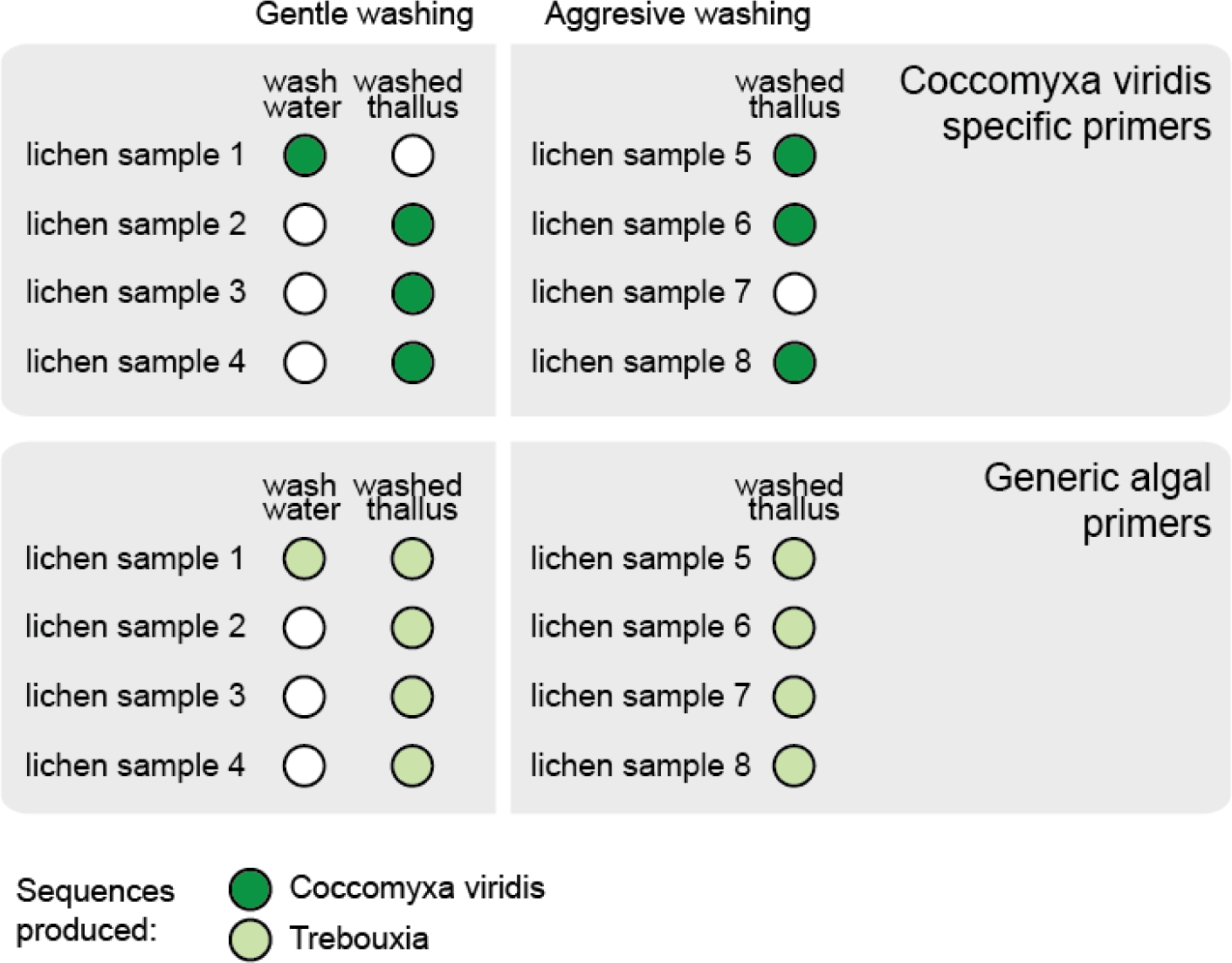
PCR-based screening for the *C. viridis* presence in washed lichen samples. Eight *X. parietina* thalli were used, of which a half were washed more gently in water, and each produced two DNA extractions: one from wash water and one from the washed thallus. The other half were washed more aggressively in ethanol and bleach; for those samples we only extracted DNA from the washed thallus. The top panel shows screening results for *C. viridis*-specific primers. Dark-green circles represent DNA extractions containing *C. viridis* DNA. Phylogenetic tree confirming the taxonomic assignment of *C. viridis* sequences is shown in Fig. S1. The bottom panel shows screening results for generic algal *rbcL* primers. Light-green circles represent DNA extractions that yielded sequences of *Trebouxia*; white circles represent DNA extractions that did not yield a usable sequence.

### Near chromosome-level genome assembly of *C. viridis* compared its relatives

Assembly of the *C. viridis* nuclear genome amounted to 45.7 Mbp and 18 contigs (Fig. 5A-B), which is close to the existing chromosome-level genome assembly from the *Coccomyxa* genus [25]. In ten contigs, we detected telomeric repeats TTTAGGG, which are typical in green algae [26]. Two contigs, cviridis_6 and cviridis_13, had telomeric sequences on both ends and likely represent complete chromosomes. The genome is estimated 97.5% complete according to BUSCO estimates and has a duplication rate of 0.2% (Fig. 5C). *De novo* annotation of the nuclear genome produced 11,248 gene models. Plastid and mitochondrial genomes were assembled in a single circular contig each (Fig. 5D-E).

**Fig. 5.**
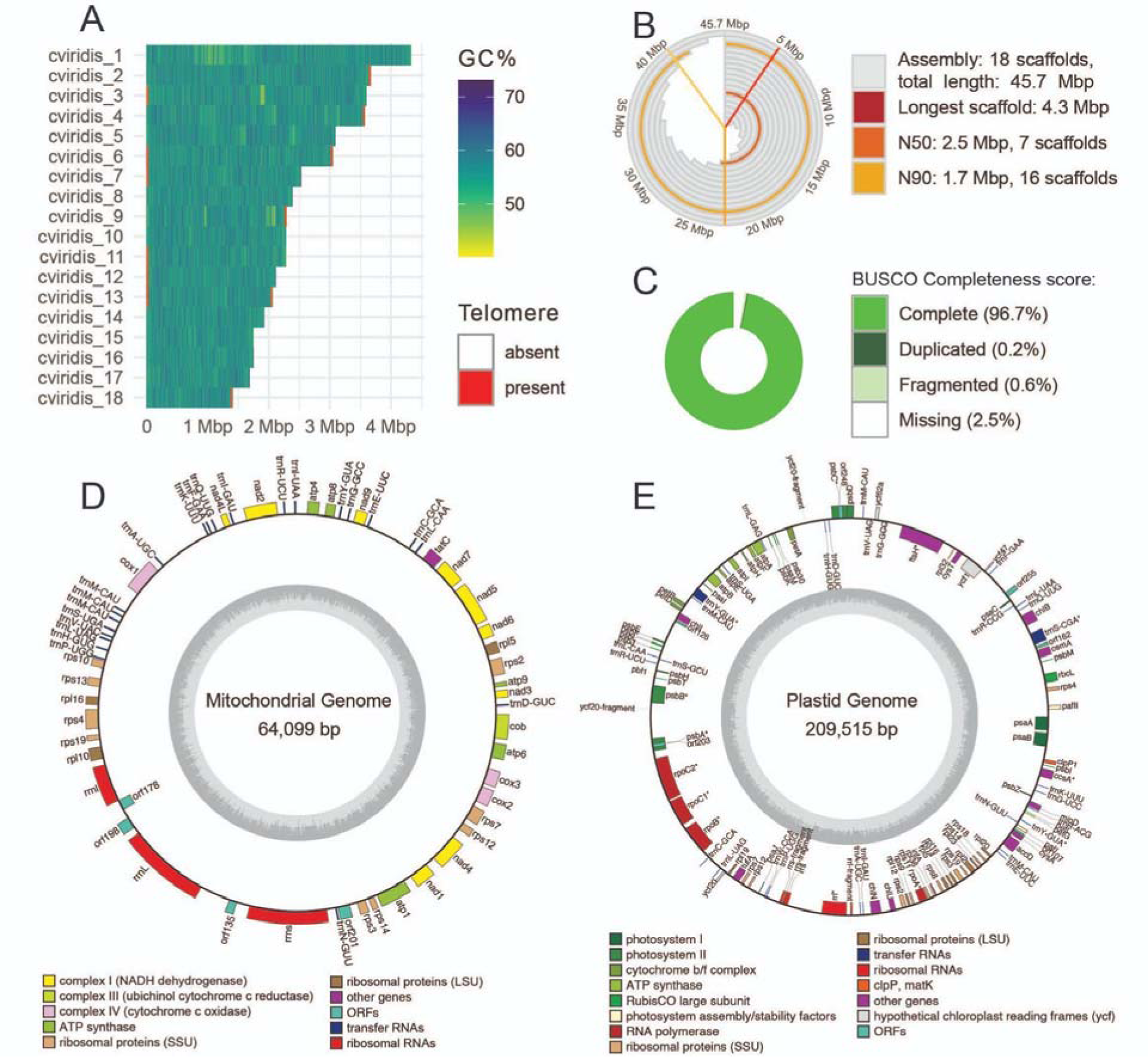
Genome of *C. viridis*. A. Plot showing GC-content and length of the 18 contigs comprising the *C. viridis* nuclear genome. Red stripes show telomeric repeats. B. Snailplot showing basic assembly statistics for the *C. viridis* nuclear genome. The gray bars show cumulative length of the assembly, with the red line showing the longest contigs. The orange and yellow lines represent N50 and N90 respectively. The plot design is based on Challis et al. [72]. C. Genome completeness scores as estimated by BUSCO (chlorophyta_odb10 database). D. Gene map of the mitochondrial genome. The inner circle represents GC-content. The genes are mapped on the outer circles, with genes inside the circle transcribed clockwise, and genes outside the circle transcribed counterclockwise. E. Gene map of the plastid genome.

In comparison to other published genomes of *Coccomyxa* algae, *C. viridis* has a slightly smaller genome, but a larger predicted proteome (Fig. 6A). The types and number of secondary metabolism gene clusters in *C. viridis* were similar to the free-living *C. subellipsoidea* (Table S3). In our functional annotations, we focused on the gene families identified by Puginier et al. [27] and Armaleo et al. [28] as connected with lichenization in green algae. Compared to other *Coccomyxa* species, *C. viridis* genome encoded comparable number of aquaporins, catalases, and domains similar to tryptophan-rich sensory protein/mitochondrial benzodiazepine receptor (TspO/MBR) (Fig. 6B) – groups of genes involved in stress response [27](Puginier et al. 2022). Unlike other studied *Coccomyxa* species, *C. viridis* genome did not encode any proteins from the Glycoside Hydrolase (GH) 8 family (as confirmed by both IntrePro and CAZy annotations) – a diverse family of hydrolases that includes licheninases, cellulases, chitosanases, and others. At the same time, it encoded one protein from the GH16 family, which also contains licheninases.

**Fig. 6.**
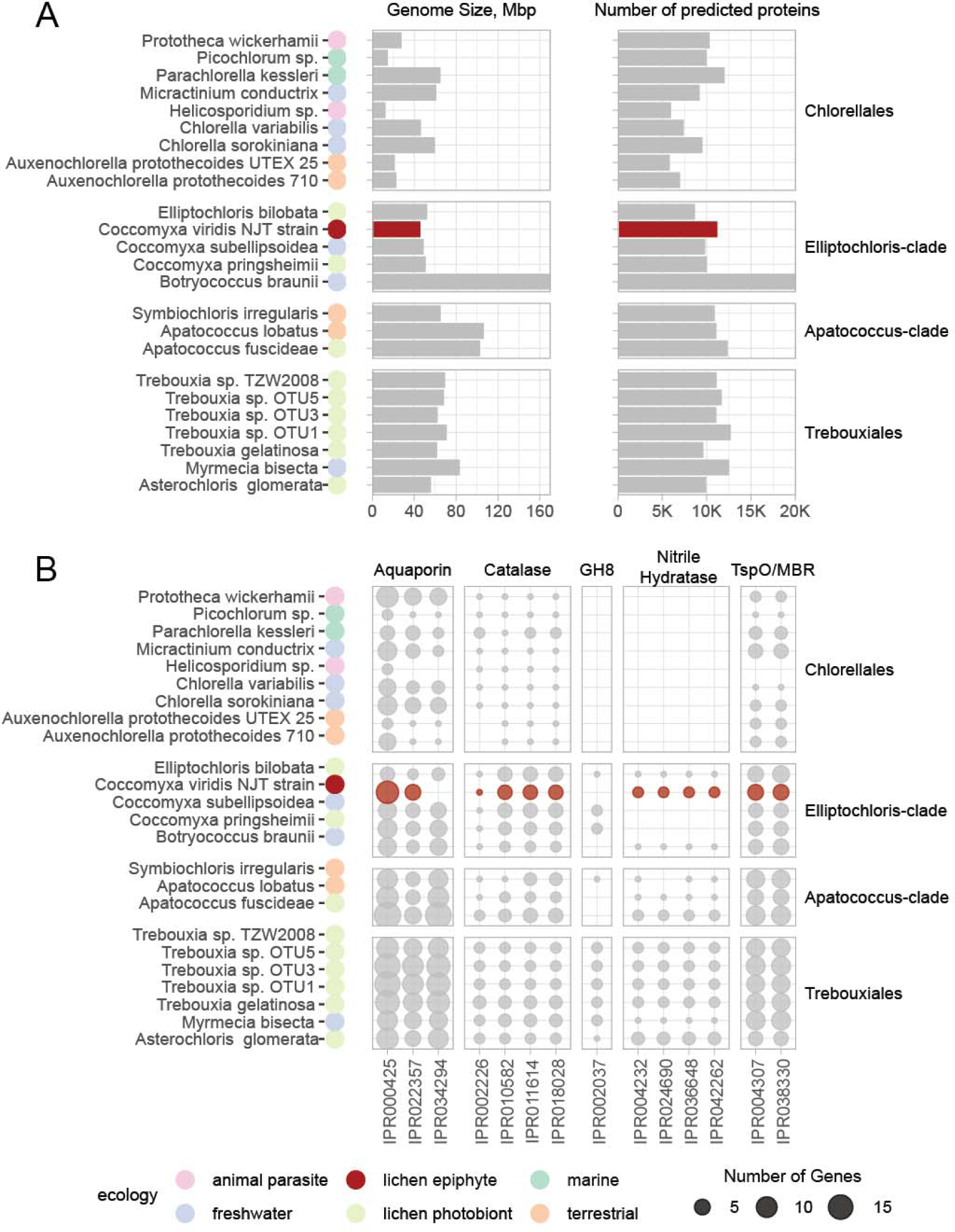
Comparative genomics analysis of *C. viridis* (highlighted in red) and other Trebouxiophyceae genomes. Information for the genomes other than *C. viridis* is taken from Puginier et al. [27]. A. Basic genome statistics plotted by taxonomic group; the circles next to the species name represent the ecology of each strain. B. Presence of InterProScan gene functional families across Trebouxiophyceae genomes. Here we show only functional families highlighted by Puginier et al. [27] as potentially relevant to lichenization in green algae. The size of the bubbles represent the number of genes assigned to each family in a given genome.

Most notably, *C. viridis* genome encoded several nitrile hydratases, which are typical in lichen photobionts [27], yet were missing from a lichen photobiont *C. pringsheimii* (Fig. 6B). The signal transduction component in *C. viridis* is largely comparable to its relatives. However, the protein kinase family (IPR000719) appears expanded in *C. viridis* (Table S4), reminiscent of similar expansions in the lichen photobiont *Asterochloris* [28].

## Discussion

Here, we present evidence that green alga *Coccomyxa viridis* is widespread in lichens as a minor component present in addition to the main photobiont. *C. viridis* has been reported before from lichens with various non-*Coccomyxa* photobionts in several isolated reports. Species from the *C. viridis* clade have been independently cultured from several lichen symbioses [13,14,16]. In addition, several amplicon metabarcoding studies of lichen algae reported small numbers of reads assigned to *C. viridis* [20,21,29]. Now, these reports are confirmed by our systematic screening of lichen metagenomic data. We detected *C. viridis* in one eighth of analyzed metagenomes. Combined with reports of other non-photobiont algae frequent in lichens [12,15], this finding raises questions about the place these algae occupy in the lichen symbiosis.

*C. viridis* comes from a genus that includes many symbiotic algae and, among others, lichen photobionts [30]. Most *Coccomyxa* photobionts come from one of the two clades: *C. subellipsoidea* and *C. simplex*/*C. solorinae,* which led to the hypothesis that lichenization happened in *Coccomyxa* twice [31]. Here, we show the *C. viridis* clade as a likely third independent origin of the lichen lifestyle, which, however, differs from the other two. In both the *C. subellipsoidea* and *C. simplex*/*C. solorinae* clades, nearly all lichen-associated algae are photobionts. Our metagenomic screening, combined with the literature data, yielded only three occurrences of *C. subellipsoidea* and *C. simplex*/*C. solorinae* in lichens with non-*Coccomyxa* photobionts (Fig. 3A, Table S1). In contrast, the *C. viridis* clade primarily contains non-photobiont algal strains isolated from lichens. Several exceptions exist, as the *C. viridis* clade includes photobionts of several *Micarea* lichens [32] and the photobiont of *Schizoxylon albescens*, an unusual lichen whose mycobiont is optionally lichenized and can occur as a non-symbiotic saprotroph [33], plus a few strains with non-lichen ecologies, including a mussel parasite [34]. Overall, the fact that *C. viridis* can occur in lichens as either a photobiont or a non-photobiont is consistent with prior reports showing ‘additional’ algae in lichen thalli to be photobionts of unrelated lichens [22]. However, our results suggest that *C. viridis*, unlike other lichen-associated *Coccomyxa* species, primarily occurs in lichens as a non-photobiont.

The newly sequenced genome of *C. viridis* is the first genome of a non-photobiont lichen-associated alga and one of the first near chromosome-level assemblies of any lichen-associated algae. It is also the third genome from *Coccomyxa*, in addition to a free-living strain of *C. subellipsoidea* [25] and the lichen photobiont *C. pringsheimii* (part of the *C. simplex*/*C. solorinae* clade) [27]. By comparing these three genomes coming from different clades and different lifestyles, we showed that they share basic genomic characteristics. At the same time, *C. viridis* exhibits more traits associated with lichenization compared to others.

What is the nature of its relationship between non-photobiont algae such as *C. viridis* and the rest of lichen symbionts? While it is possible that non-photobiont algae only treat lichens as a substrate to attach to, they can potentially reap other benefits. For lichen photobionts, participation in the symbiosis is hypothesized to bring numerous rewards: protection from herbivory, access to nitrogen, and a better hydration regime (reviewed in [35]). The extent to which non-photobiont algae have access to the same benefits might depend on whether they grow epiphytically on the surface of lichen thalli, or in the thallus interior. Our screening of washed lichen samples suggests that *C. viridis* can be endophytic, however more evidence is needed to prove this conclusively. Conversely, other lichen symbionts might benefit from the non-photobiont algae. While carbohydrates produced by a small number of *C. viridis* cells are unlikely to significantly alter the carbon budget of the lichen, the presence of a diverse set of algae could facilitate photobiont-switching thereby increasing plasticity of the symbiosis as a whole [22].

This study began with an accident. Our initial culture of the photobiont of a *Xanthoria* lichen was overgrown by *C. viridis*. Perhaps not completely coincidentally, the first sequenced genome of *Coccomyxa*, *C. subellipsoidea*, was also produced by accident in a project aimed at a different alga [25]. Relatively fast growth, observed for some non-photobiont lichen-associated algae [17], and their frequent presence make *C. viridis* contamination a likely problem in studies involving culturing of lichen symbionts. At the same time, *C. viridis* and other frequently discarded and understudied members of lichen microbiota might yet shed light on the evolution of lichen symbiosis.

Currently, we know much less about the biology and the evolution of lichen-associated algae, compared to the lichen-associated fungi. This begins to change with a recent study pioneering comparative genomics of free-living algae and lichen photobionts [27]. We believe it can be beneficial to include non-photobiont lichen-associated algae into similar studies in the future, which will now be possible with the high-quality genome of *C. viridis* we provide. As we accumulate more information on the ecology of individual algal species and in what, if any, ways they engage in lichen symbioses, we will be able to chart the evolution of lichenization in green algae.

## Methods

### Culturing

The alga was cultured from a thallus of *Xanthoria parietina* lichen kindly provided by Prof. Paul Dyer, University of Nottingham, UK. The thallus was collected in the Peak District, UK. The photobiont was isolated from the thallus as previously described [36,37]. The culture was routinely grown in liquid Bold’s Mineral Medium (BMM) on a 12-hour night/day light cycle.

### Genome sequencing and assembly

DNA was extracted from 34 mg of dry weight of algal culture, which was snap-frozen, homogenized with a Geno/Grinder homogenizer (SPEX SamplePrep, Metuchen NJ, USA) at 1,300 rpm for 1 min, and extracted with the NucleoBond High Molecular Weight DNA Kit (Macherey-Nagel, Düren, Germany). The extraction yielded 16.5 μg of high-molecular weight DNA, which was used for long-read sequencing. Short fragments were removed using Circulomics Short Read Eliminator Kit (Pacific Biosciences, Menlo Park CA, USA) with 25 kbp cut-off. A sequencing library was prepared using Native Barcoding Kit 96 V14 (Oxford Nanopore Technologies, Oxford, UK). The library was sequenced on a PromethION Flow Cell FLO-PRO114M (Oxford Nanopore Technologies, Oxford, UK) to 25 Gbp of data.

Basecalling was carried out using the ‘duplex’ method. Dorado v0.2.1 (Oxford Nanopore Technologies, Oxford, UK) was used for basecalling and Duplex tools v0.3.1 was used to identify duplex pairs. Contigs were *de novo* assembled with Flye v2.9-b1780 [38] with ‘overlap 10K, error rate 0.005, no-alt-contigs’ flags. The assembly was polished based on long reads using Medaka v1.7.2 (Oxford Nanopore Technologies, Oxford, UK). Long-read sequencing and assembly were performed by Future Genomics (Leiden, Netherlands).

In addition, we used the same DNA extraction to produce short read data. DNA was sent to Novogene UK (Cambridge, UK) and sequenced on an Illumina NovaSeq 6000 platform to 2 Gbp of PE150 data. The resulting short-read data were used to polish the long-read assembly with Pilon v1.23 [39].

### Transcriptomic sequencing

Algal culture was transferred from the liquid stock and plated on petri dishes with 99:1 BMM:MEYE culture medium. The cultures were harvested 2, 9, 21, and 42 days post inoculation, with three replicates for each time point. We snap-froze the harvested material in liquid nitrogen and extracted RNA using the RNeasy Plant Mini Kit (QIAGENE, Hilden, Germany). The RNA was sent to Novogene UK (Cambridge, UK) and sequenced on an Illumina HiSeq 2500 platform to PE150 data.

### Genome annotation

Since our initial BLASTx search against NCBI-nr showed our genomic assembly to contain bacterial sequences, we used a metagenomic binning approach to filter out contamination. We aligned Illumina reads against the assembly using Bowtie2 [40] and used the resulting bam file to bin the assembly with MetaBAT2 [41]. Next, we used the BLASTx search to select the bin that corresponded to the target algal genome. We confirmed the genome quality with BUSCO5 [42], using the chlorophyta_odb10 database. To detect telomeric repeats, we used the script from Hiltunen et al. [43] with ‘CCCTAAA’ as a query. To detect contigs representing organelle genomes, we used a BLASTx search.

Gene prediction and functional annotation of the nuclear genome was done using the Funannotate pipeline v1.8.15 [44]. We masked repeated elements in the assembly using Tantan [45] and generated gene prediction parameters using the ‘funannotate train’ command with RNA-seq data used for training. Gene prediction was performed using the ‘funannotate predict’ command, which performed *ab initio* prediction with Augustus v3.3.2 [46], CodingQuarry v2.0 [47], GlimmerHMM v3.0.4 [48], and SNAP 2006-07-28 [49]. Consensus models were created using EVidence Modeler v1.1.1 [50]. tRNA were predicted with tRNAscan-SE v2.0.9 [51]. Finally, functional annotation was done with the ‘funannotate annotate’ command. There, we assigned the gene models with putative functions based on the HMMER v3.3.2 and diamond v2.1.6 [52] searches against several databases: PFAM v35.0 [53], UniProt DB v2023_01 [54], MEROPS v12.0 [55], dbCAN v11.0 [56], and BUSCO chlorophyta_odb10 [42]. We annotated the gene models with InterPro domains using InterProScan v5.42-78.0 [57]. To annotate secondary metabolism gene clusters, we used antiSMASH v7.0.1 [58] in the fungal mode following O’Neill [59]. In order to compare our genome assembly and annotation to genomes of closely related alga, we used data from Puginier et al. [27] and Armaleo et al. [28].

Organelle genomes were annotated separately. We aligned RNA-seq data against the contigs identified as mitochondrial and plastid genomes using STAR v2.5.4b [60] and predicted genes using MFannot [61] and GeSeq [62]. To finalize the annotations, we manually combined the outputs of the two tools and cross-referenced it against the RNA-seq alignment. The annotations were visualized using the OGDraw webserver [63].

### Phylogenetic analyses

To provide a taxonomic identification to the sequenced genome, we first built a phylogenomic tree using 10 reference genomes and transcriptomes from Trebouxiophyceae with *Clamydomonas eustigma* as an outgroup (Table S5). We identified chlorophyta_odb10 BUSCO single-copy orthologs shared by all genomes and transcriptomes, which amounted to 196 loci. Next, we created a single concatenated alignment using MAFFT v7.271 [64] and trimmed it with trimAL v1.2 [65] to remove positions present in <70% of organisms. Finally, we computed a phylogeny with RAxML v8.2.12 [66], using PROTGAMMAAUTO model. To provide a better taxonomic resolution, we created a tree based on ITS sequences. We included 77 reference ITS sequences from *Coccomyxa* and *Elliptochloris* (Table S6). The tree was constructed as described above.

### Screening of publicly available metagenomic data

We searched for *Coccomyxa viridis* ITS in the 471 metagenomic assemblies from Tagirdzhanova et al. [67] using a BLASTn search with the e-value cut-off of 1e-65 (Table S7). As a query, we used the ITS sequence pulled from the genome assembly. Extracted hits were combined with the ITS reference sequences (Table S2), aligned as described above, and used to construct a phylogeny using IQTREE v2.2.2.2 [68].

### Screening of lichen thalli

To determine if *Coccomyxa viridis* is external or internal to lichen thalli, we screened eight thalli of *Xanthoria parietina* lichen, in part following Moya et al. [18]. The lichen samples were collected in Norwich Research Park (Norwich, UK; 52.623133°N, 1.221621°E) from tree bark. We separated a 1 cm^2^ fragment of each thallus from its substrate taking care to remove all visible fragments of bark, moss, or other contaminants.

Four fragments were subjected to ‘soft’ washing. We soaked them in filter-sterilized water for 10 min, then vortexed for 5 min at 600 rpm. Next, we brushed the upper surface of thallus fragments with a soft paintbrush, following Yoshimura et al. [69] and washed it in a jet of deionized water. Each thallus fragment yielded two samples: (1) washed lichen fragment and (2) wash water sample. We centrifuged the wash water samples at 4,000 rpm for 5 min to obtain cell pellets.

Next, we dried both washed lichen fragments and wash-water pellets at 65°C and extracted DNA with the DNeasy Plant Mini Kit (QIAGEN, Hilden, Germany).

The second half of fragments were washed more aggressively. We followed the protocol from U’ren et al. [70] for surface-sterilization of lichen thalli. First, we washed each fragment in a jet of deionized water. Next, we soaked and vortexed fragments at 600 rpm in three solutions: 95% ethanol for 10 sec, 0.525% NaOCl for 2 min, and 75% ethanol for 2 min. Finally, the thalli were washed in a jet of deionized water for 2 min. We extracted DNA from the thalli as described above.

We screened the DNA extractions using two pairs of primers: specific for *Coccomyxa viridis* ITS, and general green algal *rbcL* primers (Table S8). PCR reactions were performed using Q5 High-Fidelity DNA Polymerase (New England Biolabs, Ipswich MA, USA). For ITS we used the following conditions: 98°C for 5 min, followed by 50 cycles of 98°C for 30 sec, 70°C for 30 sec, and 72°C for 30 sec, followed by the final extension step of 72°C for 7 min. For *rbcL* we used the following conditions: 98°C for 5 min, followed by 35 cycles of 98°C for 1 min, 57°C for 1 min, and 72°C for 1 min, followed by the final extension step of 72°C for 7 min. PCR reactions were gel-extracted using the Wizard SV Gel and PCR Clean-Up System (Promega, Madison WI, USA) and sequenced by GENEWIZ (Leipzig, Germany). We used the resulting sequences to build a phylogeny with reference sequences as described above.

### Microscopy

We visualized the culture of C. viridis and, for comparison, a culture of a *Trebouxia* photobiont isolated from a *X. parietina* thallus (the thallus was collected in Norwich Research Park; *Trebouxia* was isolated as described above). Algal samples were stained in Calcofluor White Stain (Sigma-Aldrich, Burlington MA, USA) for three minutes. Confocal microscopy was performed using a Leica SP8 laser confocal microscope with excitation wavelength 405 nm and emission wavelengths 410-430 nm for calcofluor white and 650-730 nm for the chlorophyll autofluorescence. In addition, we performed bright-field imaging of a cross-section through a *X. parietina* thallus using a Leica DM5500b microscope. Images were analyzed with Leica software and Fiji.

## Acknowledgements

This work was supported by grants from the Leverhulme Trust RPG-2018-139, The Gatsby Charitable Foundation, the Halpin Family and the Biotechnology and Biological Sciences Research Council BBS/E/J/000PR9798 to N.J.T. The authors thank Paul Dyer and Peter Crittenden for providing the algal culture. The authors thank Camille Puginier, Francesco Dal Grande, and other authors of Puginier et al. [27] for sharing with us genome statistics behind Fig. 1 and Fig. 2 of their study. We thank Pavel Škaloud for discussions, and Phil Robinson and Jason Hollinger for providing photographs.

## Author contributions

G.T. and N.J.T. designed the study. G.T., K.S., and X.Y. performed experimental work. G.T. performed bioinformatic analysis. X.Y. and G.T. performed microscopy. G.T. and N.J.T. drafted the manuscript, and all authors contributed to editing.

## Data and Code Availability

The annotated genome assembly of *C. viridis* will be available at NCBI (Study Accession PRJEB65893; release pending). All code associated with the analysis along with details on the usage of bioinformatics tools is available on GitHub (https://github.com/metalichen/2023_Coccomyxa_viridis_genome).

