## Supplementary Figures for "Genomic analysis of *Coccomyxa viridis*, a common low-abundance alga associated with lichen symbioses"

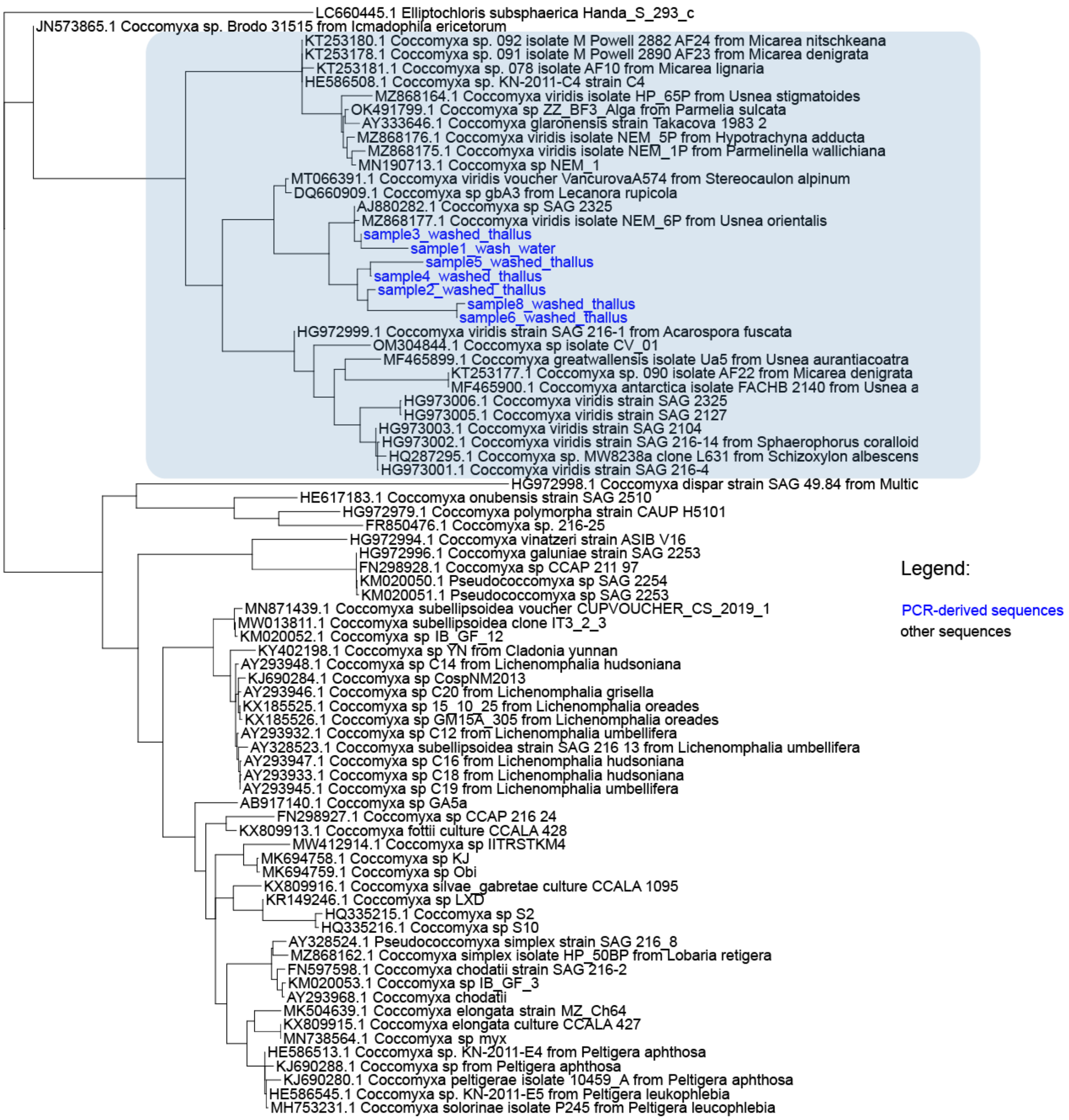


Fig. S1. Maximum-likelihood phylogenetic tree of *Coccomyxa* ITS. Sequences produced by PCR highlighted in blue font. The blue rectangle shows the *C. viridis* clade.
